# Link clustering explains non-central essential genes in protein interaction networks

**DOI:** 10.1101/532770

**Authors:** Inhae Kim, Heetak Lee, Seong Kyu Han, Kwanghwan Lee, Sanguk Kim

**Affiliations:** Department of Life Sciences, Pohang University of Science and Technology, Pohang 790-784, Korea; School of Interdisciplinary Bioscience and Bioengineering, Pohang University of Science and Technology, Pohang 790-784, Korea

**Author notes:** Correspondence should be addressed to: Sanguk Kim, Ph.D., Department of Life Sciences, Pohang University of Science and Technology, Pohang 790-784, Korea.

## Abstract

Essential genes (EGs) often form central nodes in protein-protein interaction (PPI) networks. However, many reports have shown that numerous EGs are non-central, suggesting that another principle governs gene essentiality. We propose link clustering as a distinct indicator of the essentiality for non-central nodes. Specifically, in various human and yeast PPI networks, we found that 29 to 47% of EGs were better characterized by link clustering than by centrality. Such non-central EGs with clustered links have significant impacts on communities at lower hierarchical levels, suggesting that their essentiality derives from functional dependency among relevant local neighbors, rather than their implication on global connectivity. Moreover, these non-central EGs exhibited several distinct characteristics: they tend to be younger and fast-evolving, and likely change their essentiality across different human cell lines and between human and mouse than central-EGs.

**Author summary:** The centrality in network structure was thought to be the cause of gene essentiality, as it conveys nodes’ importance in given networks. However, many essential genes are also found to be non-central, which leads us to ask whether a principle other than centrality may govern gene essentiality. Here we demonstrate that link clustering conveying functional dependency between nodes is the other principle characterizing gene essentiality. The link clustering explains numerous essential genes that are non-central, which are distinct from central ones regarding their molecular function and evolution. Our results clearly establish a two-dimensional principle governing gene essentiality.

## Introduction

The centrality-lethality (C-L) rule states that central genes in PPI networks are likely essential for the survival and the reproduction of the organism [1]. Specifically, the number of partners, or the degree, in a PPI network directly correlates with a gene’s essentiality. The rule is attributed to “attack vulnerability” of networks exhibiting scale-free degree distribution, in which the removal of the most connected nodes rapidly results in the loss of global network connectivity [2]. Therefore, the C-L rule proposes that centrality is the cause of gene essentiality, as it conveys the functional importance of central genes in network structure. Other centrality measures, which defined “bottleneck” and “minimal determining set”, further supported the C-L rule as a complementary notion to the degree [3–5].

However, it is questionable whether a node’s capability to sustain global connectivity is the sole cause of gene essentiality. Importantly, the C-L rule itself suggests that a considerable amount of non-central genes may be essential. The key idea underlying the C-L rule is that scale-free networks exhibit “error tolerance” at the cost of “attack vulnerability” [1, 2]. These networks are tolerant to random failure (i.e., error) of a node, since they contain an extremely large number of non-central genes that have only a slight chance to be essential. Therefore, by the C-L rule, centrality predicts not only the chance of being essential, but also the number of genes with given chance: the smaller the chance a gene has to be essential, the greater the frequency such genes are in the network. By the virtue of frequency, the expected amount of non-central EGs is not necessarily negligible. Interestingly, recent studies showed that non-central EGs may convey a distinct biological relevance. For instance, yeast EGs that were evolvable (i.e., became non-essential) under a laboratory condition showed smaller interaction degree than non-evolvable ones [6]. In addition, cell line-specific EGs across different human cells exhibited smaller degree than genes that are universally essential [7]. This begs the question of whether there exists another principle governing gene essentiality that is orthogonal to network centrality.

Here we hypothesize that highly clustered links contribute to gene essentiality, as such links represent functional dependency between nodes. It was previously proposed that links with stronger functional dependency have a greater impact on network robustness, as the failure of one node will likely result in the failure of its neighborhood [8, 9]. At molecular level, such an example among proteins is obligate interactions: a protein is unstable on its own, thus has to be bound its partner to sustain its stability [10]. Interestingly, Pajevic and Plenz showed that link clustering from network structure can estimate empirically observed functional dependency in many networks, such as gene co-expression and regional co-activation in the cerebral cortex [11]. Moreover, link clustering may characterize functional importance of a gene independently of centrality measures. Link clustering estimates functional dependency by gauging neighborhood overlap between two interacting nodes[11–13] (Fig 1A), and central nodes likely convey low functional dependency for their many non-overlapping interactions (Fig 1B). Indeed, the link clustering showed a distinct implication on networks’ global and local connectivity compared with centrality [11].

**Fig 1.**
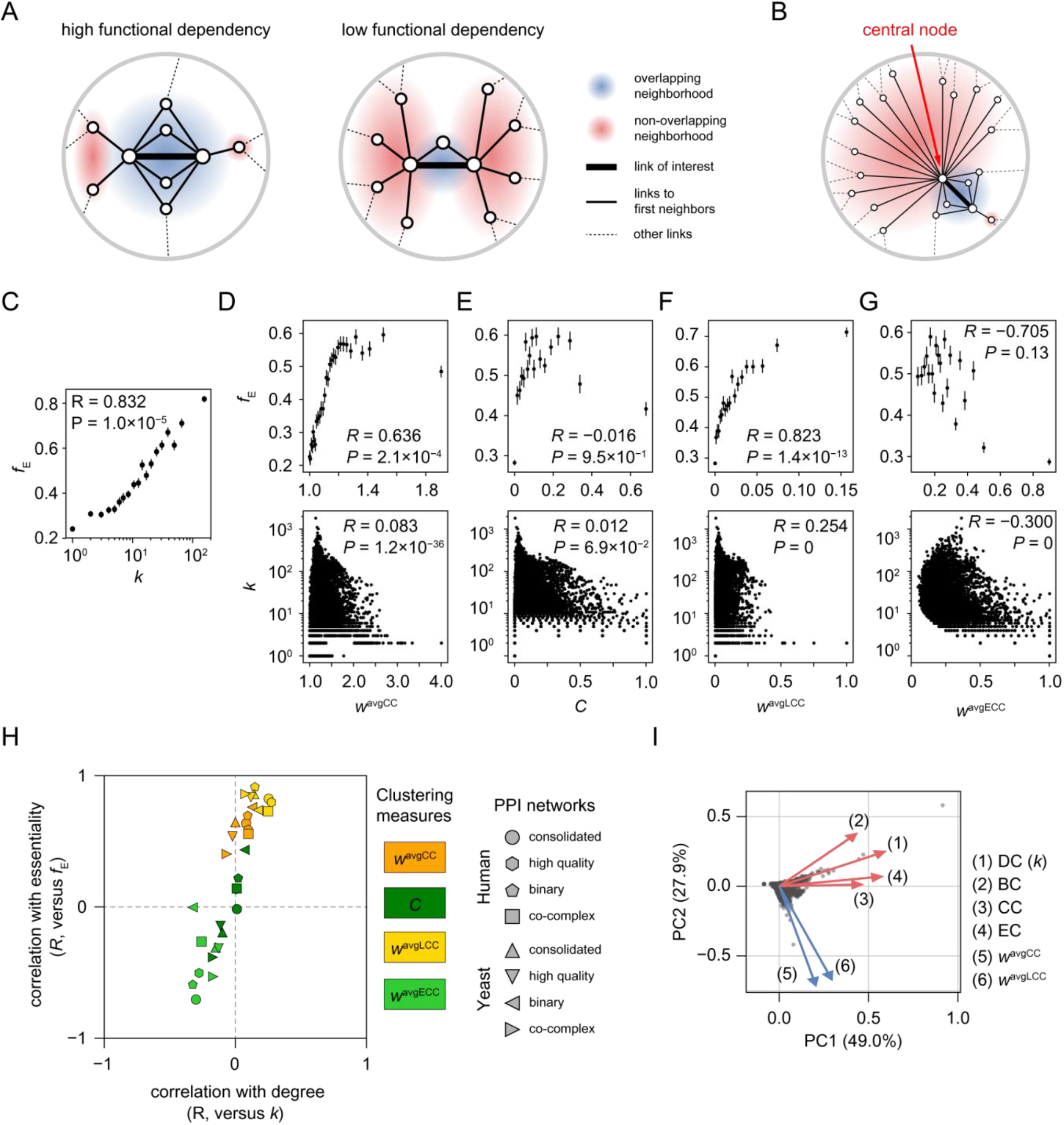
The relationship between gene essentiality and centrality and clustering measures in PPI networks. (A) Link clustering estimates functional dependency by measuring neighborhood overlap. (B) A central node likely conveys low functional dependency for its many non-overlapping neighbors. (C) Correlation between the degree (k) and the fraction of gene essentiality fE), which confirms the C-L rule. To explore different clustering measures, scatter plots were constructed for (D) *w*^avgCC^, (E) C, (F) *w*^avgLCC^, and (G) *w*^avgECC^ against *f*_E_ (upper panel) and *k* (lower panel). Error bars of *f*_E_ indicate the standard deviation of the normal distribution approximated from binomial trials with success probability estimated by the fraction of EGs. (H) Summary of the observed correlation between clustering measures and *f*_E_ and *k* in eight different PPI networks. (I) PCA analysis of centrality and link clustering measures on EGs. Number in parenthesis indicates the variance explained by the component.

Notably, previous studies showed that EGs are relevant to network clustering: they tend to be directly connected [14, 15] or enriched within same functional modules and protein complexes [16–21]. Unfortunately, such clustered EGs seem to be central, rather than featuring non-central EGs. It was shown that EGs within essential modules exhibit greater degree in PPI networks than those within non-essential modules [18], and interactions between EGs were more frequently found between genes with greater degree [14]. In addition, essential modules have greater interaction degree in module-level networks [22]. These observations strongly suggest that EGs are often clustered *and* central. With respect to network structure, this is somewhat expected, as central genes are frequently clustered together due to their greater connectivity [23]. Therefore, toward our goal for understanding non-central EGs, clustered network structure has not yet been sufficiently characterized.

In this study, we aimed to identify non-central EGs associated with clustered network structure. We systematically compared various cluster measurements for their ability to characterize node essentiality and orthogonality to centrality. We found that link clustering is an accurate indicator of node essentiality independent of centrality, enabling us to correctly classify a substantial number of non-central EGs. These EGs with clustered links have a significant impact on small communities with a low-level hierarchy, suggesting that their essentiality derives from functional dependency among relevant local neighbors. Additionally, these EGs also exhibit distinct evolutionary features, as they are younger and evolve faster, and tend to be conserved as a group with neighboring genes. Moreover, in human, they likely change their essentiality across cell lines and species.

## Results

### Link clustering is a key network topological parameter to characterize non-central EGs

To analyze the relationship between clustered links and gene essentiality, we defined a link weight, proportional to the clustering coefficients of two end nodes, similar to that proposed by Pajevic and Plenz [11]:

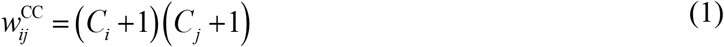

where *C* represents the node clustering coefficient, calculated as *C_i_* = 2*T_i_*/(*k_i_* × [*k_i_* − 1]), in which *T_i_* represents the number of triangles including node *i*, and *k_i_* represents the number of interaction partners, or the degree, of node *i*. We added one to each node clustering coefficient to avoid the case in which *C_i_* = 0 nullifies *C_j_* ≠ 0. A node’s association with its local neighbors is then quantified by taking the average weight of the connected links:

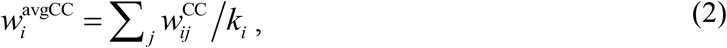

where node *j* indicates an interaction partner of node *i*.

As the C-L rule anticipates, in a human integrated PPI network (dubbed as “consolidated”), we found that the fraction of EGs, *f*_E_, increases as a function of *k* (Fig 1C, Pearson correlation coefficient *R* = 0.832, *P* = 1.0×10^−5^). Additionally, *f*_E_ increases as a function of *w*^avgCC^ (Fig 1D, upper panel; *R* = 0.636, *P* = 2.1 ×10^−4^), demonstrating that clustered links are also related to gene essentiality as we hypothesized. We observed a poor correlation between *w*^avgCC^ and *k* (Fig 1D, lower panel; *R* = 0.083, *P* = 1.2×10^−36^), indicating that genes with large *w*^avgCC^ are unlikely to be central in the network although many of them are essential. Therefore, this observation suggests that *w*^avgCC^ may provide an explanation for EGs that are non-central.

Unlike *w*^avgCC^, the node clustering coefficient (*C*) was poorly correlated with both *f*_E_ (Fig 1E, upper panel; *R* = −0.016, *P* = 0.95) and *k* (lower panel; *R* = 0.012, *P* = 0.069). This suggests that *w*^avgCC^ and *f*_E_ are positively correlated because of the partners’ clustering coefficients. To assess the interaction partners’ contribution to the relationship between *w*^avgCC^ and *f*_E_, we combine equations (1) and (2),

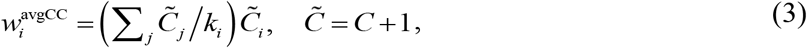

where node *j* represents an interaction partner of node *i*. This equation shows that the slope of a straight line between 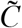 and *w*^avgCC^ quantifies the partner’s contribution. With least square fits of EGs and non-EGs, we found that the slope for EGs (*β*_E_ = 1.50) was significantly greater than that for non-EGs (S1 Fig, *β*_NE_ = 1.32, *P* = 1.9×10^−86^). This result emphasizes that clustered links rather than clustered nodes are crucial for characterizing EGs.

### Functional dependency of link clustering can explain gene essentiality

Given that node clustering itself is not informative for gene essentiality, we asked whether the functional dependency represented by link clustering is relevant to gene essentiality. Specifically, we investigated two more clustering measures gauging the extent of neighborhood overlap between end nodes. One measure used was the link clustering coefficient, *w*^LCC^, previously shown to correlate with link strength characterizing co-functioning of the end nodes [11, 13] and defined by the equation:

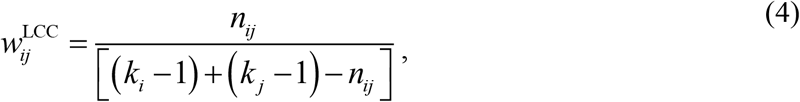

where *n_ij_* is the number of common neighbors of node *i* and *j*. The other measure used was the edge clustering coefficient, *w*^ECC^, devised to estimate the extent of the end nodes being within a same community [12] and defined by the equation:

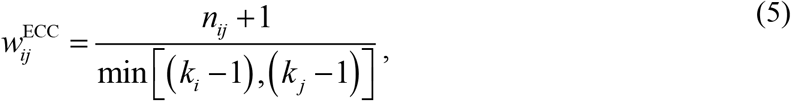

where min [(*k_i_* − 1),(*k_j_* − 1)] is the maximal possible number of common neighbors. We then summarized link weights onto nodes by taking the average; *w_i_* = Σ_*j*_ *w_ij_* / *k_i_*.

We found that only *w*^avgLCC^ is a good indicator of gene essentiality although it did not have correlation with centrality. Specifically, *w*^avgLCC^ positively correlated with *f*_E_ (Fig 1F, upper panel; *R* = 0.823, *P* = 1.5×10^−5^) but did not with *k* (lower panel; *R* = 0.254, *P* = 0), similar to *w*^avgCC^ (Fig 1D). In contrast, *w*^avgECC^ negatively correlated with both *f*_E_ (Fig 1G, upper panel; *R* = −0.705, *P* = 3.6×10^−4^) and *k* (lower panel; *R* = −0.300, *P* = 0). The fundamental difference between these clustering measures is how they account for two end nodes of a link. Whereas *w*^ECC^ represents one node’s dependency to the other by normalizing neighborhood overlap over only one node’s degree, *w*^LCC^ characterizes the dependency between two nodes by considering both degrees. Therefore, it seems crucial to quantify functional dependency of a link by considering both end nodes when one attempts to characterize gene essentiality.

We further examined whether these link clustering measures from network structure are relevant to the functional dependency between protein molecules in interaction proteomics studies [24–27]. Specifically, these experiments provide link weight (*w*^EXP^), which quantifies the reliability in the assembly of protein molecules, representing the functional dependency of protein interactions. Taking *w_i_* = Σ_*j*_ *w_ij_* / *k_i_*, we observed that both *w*^avgCC^ (S2A Fig) and *w*^avgLCC^ (S2B Fig) positively correlated with *w*^avgEXP^, indicating that the link clustering measures are indeed good estimates of functional dependency. In addition, we observed a strong positive correlation between *w*^avgEXP^ and *f*_E_ in all experiments (S2C Fig; S1 Table), confirming that functional dependency between proteins explains gene essentiality.

We confirmed the robustness of these results in various PPI networks from two different eukaryotic organisms – human and yeast. For each species, we tested four different PPI networks, including “consolidated,” “high-quality,” “binary,” and “co-complex” networks, as it was previously reported that the stringency of quality control and the variations in PPI detection methods affect network structure [28] (S1 Table; see Discussion). We found that two important observations remained unchanged in the analyses of different PPI networks (Fig 1H). First, none of the four clustering measures positively correlated with *k* (Fig 1H, horizontal axis). Second, both *w*^avgCC^ (Fig 1H, orange markers) and *w*^avgLCC^ (yellow) positively correlated with *f*_E_, whereas *C* (green) and *w*^avgECC^ (light green) showed a weak correlation (vertical axis). These results were robust with other centrality measures, including betweenness (*BC*), closeness (*CC*), and eigenvector centrality (*EC*) (S3 Fig and S2 Table). Accordingly, principal component analysis on EGs showed that the clustering measures were separated from the centrality measures, while the two components explained a large fraction of the whole variance among EGs (Fig 1I; S4 Fig for all PPI networks). Additionally, we confirmed the partners’ contributions to gene essentiality using equation (3), as the slope of EGs in the linear fit (*β*_E_) was significantly greater than that of non-EGs (*β*_NE_) in all 8 PPI networks (S1 Fig). Taken together, these results strongly indicate that link clustering explains the essentiality of non-central genes due to the functional dependency it conveys, rather than the clustered network structure itself.

### Centrality and link clustering have distinct implications on network connectivity

We further examined that centrality and clustering measures depict inherently distinct properties of network structure in terms of global and local connectivity. For simplicity, we first focus on two measures, *k* and *w* = *w*^avgCC^, and later expand the analysis to include other measures. We consecutively removed nodes with the greatest *k* (*k*-pruning) or *w* (*w*-pruning) values and monitored how robust the network was to node removal in terms of global or local connectivity. Connectivity is characterized by two distinct parameters, the relative size of the giant component, *R*_GC_, and the excess clustering normalized by the degree sequence preserved randomization, Δ*C*. While *R*_GC_(*f*) can be used to represent global connectivity by quantifying the fraction of all accessible nodes in the largest connected component, Δ*C*(*f*) summarizes local connectivity by measuring the average fraction of links between only the first neighbors of individual nodes, where *f* represents the fraction of removed nodes.

We found that *k* has a greater impact on the global connectivity, while *w* has a greater impact on local connectivity. In the human consolidated network, for example, *R*_GC_(*f*) decreased more rapidly in response to *k*-pruning than to *w*-pruning (Fig 2A), while Δ*C*(*f*) decreased more rapidly in response to *w*-pruning than to *k*-pruning (Fig 2B). We then compared the effect of random pruning (r-pruning), during which nodes are removed in a random order. With *r*-pruning, *R*_GC_(*f*) and *f* exhibited a straight line (Fig 2A and 2B; grey dotted line), indicating that random removal does not have an excessive impact on global connectivity.

**Fig 2.**
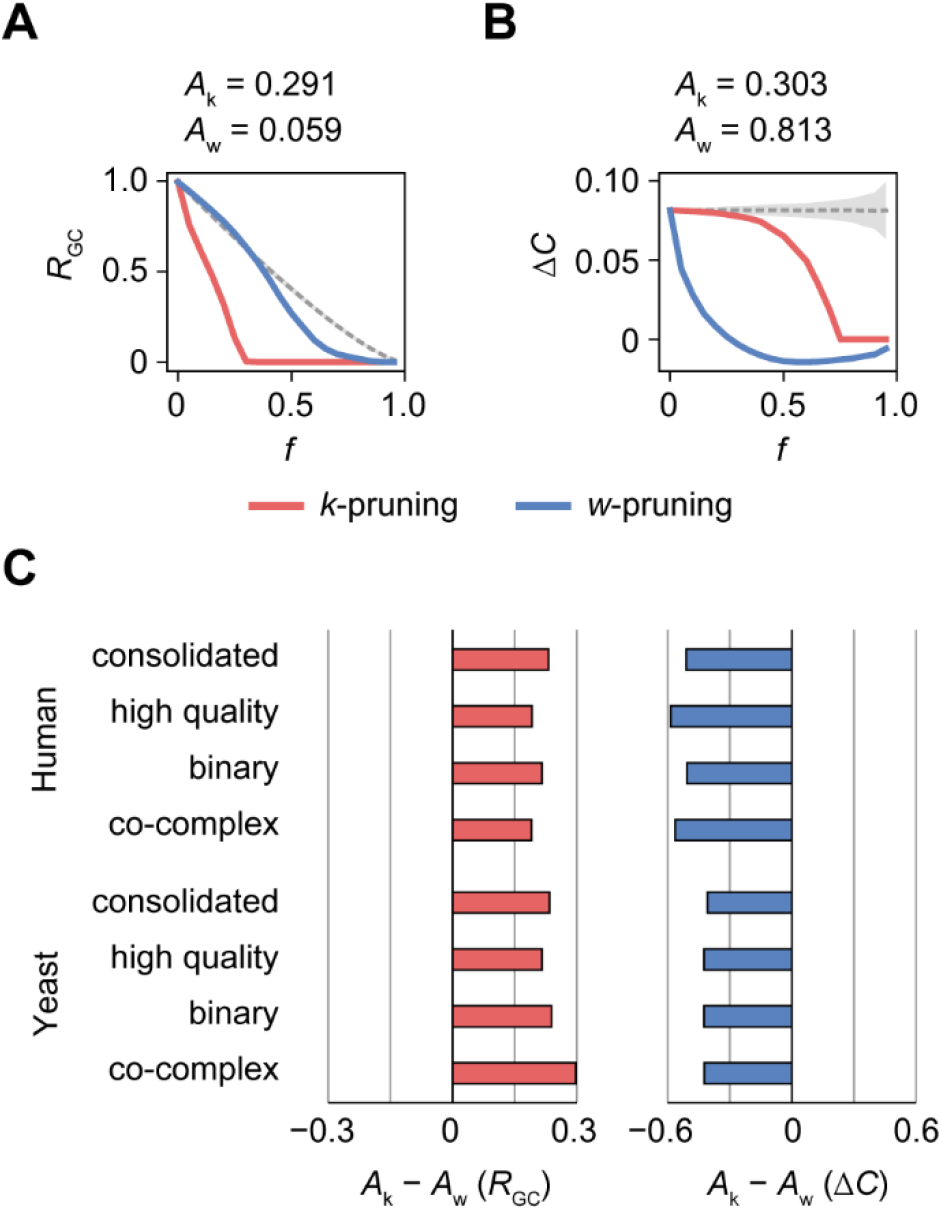
Comparison of the impacts of *k* and *w* on network connectivity. (A) The change in global connectivity (*R*_GC_) with node removal in decreasing order of *k* (red lines) and *w* (blue lines), or in random order (grey dotted lines; area, ±3*σ*) with the fraction, *f*, of removed nodes. (B) The change in local connectivity (Δ*C*) with node removal. (C) Differences in *R*_GC_ and Δ*C* with *k*- and *w*-pruning quantified as the area under the random pruning curve; *A*_k_ represents the difference in area under the curve between the *k*- and *r*-pruning curves, and *A*_w_ represents the difference in area under the curve between the *w*- and *r*-pruning curves. To calculate the area, *R*_GC_ and Δ*C* were normalized to a range from 0 to 1.

To quantify these differences, we compared the area between the *k*- and *r*-pruning curves and the area between the *w*- and *r*-pruning curves (see the Methods section). Specifically, we measured the area over the *k*- or *w*-pruning curve and under the *r*-pruning curve; the area was negative when the *k*- or *w*-pruning curve was placed over the *r*-pruning curve. For *R*_GC_(*f*), the area of the *k*-pruning curve (Fig 2A; *Ak* = 0.291) was greater than that of the *w*-pruning curve (*A*_w_ = 0.059), indicating that global connectivity is more vulnerable to the removal of nodes with greater *k* values. By contrast, Δ*C*(*f*) decreased more rapidly with *w*-pruning than with *k*-pruning (Fig 2B; *A*_k_ = 0.303, *A*_w_ = 0.813), demonstrating that local connectivity is more vulnerable to the removal of nodes with greater *w* values. We observed similar results in different PPI networks: *A*_k_ - *A*_w_ > 0 for *R*_GC_(*f*) and *A*_k_ - *A*_w_ < 0 for Δ*C*(*f*) (Fig 2C). We also confirmed that the result is consistent with other centrality measures. In all 32 cases (4 centrality measures × 8 networks), *R*_GC_(*f*) decreased more rapidly by centrality measures and Δ*C*(*f*) decreased more rapidly by *w*^avgCC^ (S5A and S6 Fig).

We then investigated pruning by *w*^avgLCC^, which also proved to characterize gene essentiality distinctively from centrality measures. In general, *w*^avgLCC^ behaved similarly to *w*^avgCC^: *R*_GC_(*f*) decreased more rapidly by centrality measures and Δ*C*(*f*) decreased more rapidly by *w*^avgLCC^ (S5B and S7). However, the decrease of Δ*C*(*f*) by *w*^avgLCC^ was quite similar to that achieved by centrality measures in several yeast networks (S7 Fig). This indicates that, compared to *w*^avgLCC^, *w*^avgCC^ is more distinct from centrality measures. Therefore, we chose *k* and *w* = *w*^avgCC^ to represent centrality and clustering, respectively, throughout the rest of the manuscript unless otherwise stated.

### Clustered links characterize a distinct set of EGs

Given that gene essentiality is characterized by two distinct properties, *k* and *w*, we may expect that EGs are composed of two different subsets: those better characterized by *k* (*k*-dependent), and those better characterized by *w* (*w*-dependent). Using logistic regression, we calculated the probability of a gene being essential based on *k*, *P*_E_(*k*), and based on *w*, *P*_E_(*w*), and then classified the genes as *k*-dependent when *P*_E_(*k*) > *P*_E_(*w*) or as *w*-dependent when *P*_E_(*k*) < *P*_E_(*w*) (S8 Fig). We also considered a third case, in which EGs are explained neither by *k* nor by *w*. Specifically, we determined kc and wc as the cutoffs to maximize Matthew’s correlation coefficient (MCC) by regarding genes with *k* ≥ k_c_ or *w* ≥ w_c_ as predictive EGs, and discarded genes satisfying *k* < k_c_ or *w* < w_c_ from the *k*- or *w*-dependent sets, respectively (S9 Fig).

In our data set, a sizable number of EGs were *w*-dependent. The criteria used to classify a gene as a *k*- or *w*-dependent essential gene, as well as the *P*_E_(*k*) and *P*_E_(*w*) values of the EGs in the human consolidated network, were shown in Fig 3A. We found that 36.0% of EGs were *w*-dependent (*n* = 2,186; Fig 3A, dark blue and pale blue circles), a proportion comparable to that of *k*-dependent EGs (40.9%, *n* = 2,483; dark red and pale red circles). In the eight different PPI networks, 29 to 47% of EGs were *w*-dependent (Fig 3B). It is also notable that many *w*-dependent EGs had *k* < k_c_ (*n* = 1,570, dark blue circles; Fig 3A), indicating that they are unlikely classified as essential by centrality. In the eight different PPI networks, 10 to 25% of the EGs were *w*-dependent with *k* < k_c_ (Fig 3B). All *k*- and *w*-dependent EGs in yeast and human PPI networks were shown in S3 and S4 Table, respectively.

**Fig 3.**
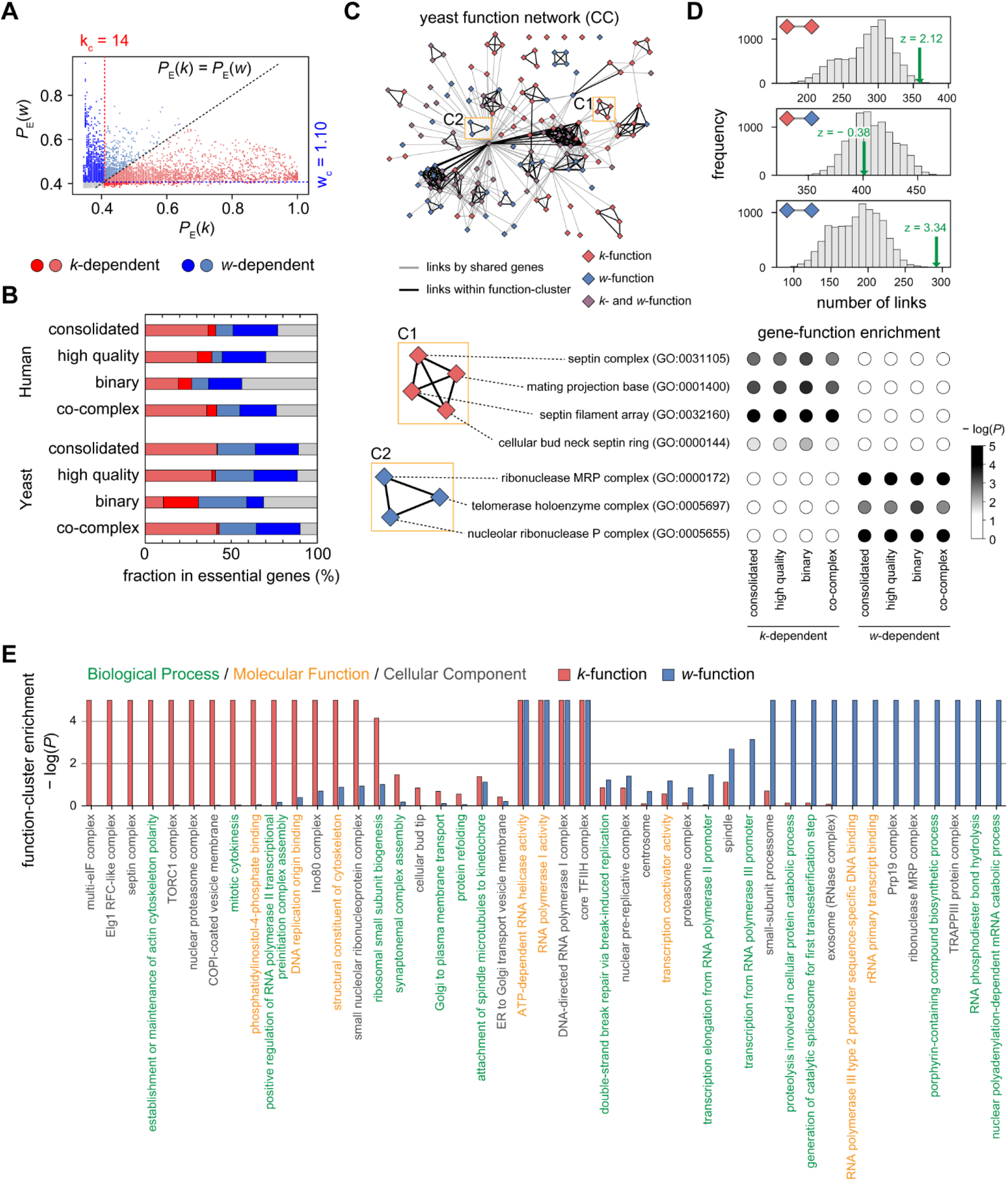
Classification of *k*- and *w*-dependent EGs and their functional differences. (A) Probability of being essential based on k and *w* of all EGs in the human consolidated network. (B) Fraction of *k*- and *w*-dependent EGs in the eight different PPI networks examined. (C) A functional network of yeast cellular component (CC) terms connected by shared genes. Terms were identified as *k*- or *w*-functions by their enrichment with *k*- and *w*-dependent EGs from three or four PPI networks. Two clusters, C1 and C2, are shown as examples. (D) Number of links between *k*- and *w*-functions in the functional network of yeast CC terms; upper panel, between two *k*-functions; middle panel, between *k*- and *w*-function; lower panel, between two *w*-functions. Green arrows indicate the number observed in the real network, and grey bars show the distribution of the numbers observed from 10,000 random sets with shuffled *k*- and *w*-function tags. (E) Representative GO terms of clusters from yeast functional networks and the bias of clusters towards *k*- and *w*-functions.

We next examined whether *k*- and *w*-dependent EGs are distinct not only with respect to network structure but also biological function (see the Methods section). To provide a comprehensive comparison of biological function, we defined *k*- and *w*-functions, which are gene ontology (GO) terms enriched with *k*- and *w*-dependent EGs, in consensus with three or four PPI networks. We then constructed functional networks of GO terms connected by shared genes, and using these functional networks, we investigated clustered *k*- and *w*-functions.

We found that *k*- and *w*-dependent EGs in our networks were often separately associated with distinct biological functions, which were partitioned in the functional networks. For example, in the yeast functional network comprised of Cellular Components (CC) terms, *k*- and *w*-functions emerged together into a large and well-connected network (Fig 3C; S10 Fig). However, many clusters were biased towards *k*- or *w*-functions. For instance, one cluster included four similar *k*-functions, “septin complex,” “mating projection base,” “septin filament array,” and “cellular bud neck septin ring,” that were enriched with *k*-dependent EGs from all four PPI networks (Fig 3C; box C1). Another cluster possessed three related *w*-functions, “ribonuclease MRP complex,” “telomerase holoenzyme complex,” and “nucleolar ribonuclease P complex,” that were enriched with *w*-dependent EGs from all four PPI networks (Fig 3C; box C2). Links were observed more frequently between two *k*-functions (Fig 3D, upper panel; *n* = 359, *z* = 2.12) and between two *w*-functions (Fig 3D, lower panel; *n* = 292, *z* = 3.34) compared with 10,000 random sets with shuffled *k*- and *w*-function tags. In contrast, links between *k*- and *w*-functions were observed similar to random expectation (Fig 3D, middle panel; *n* = 401, *z* = −0.38). This result was robust over all functional networks comprised of three GO categories in yeast and human (S11 Fig).

This approach enabled us to complete a comprehensive summary of biological functions associated with *k*- and *w*-dependent EGs (Fig 3E, yeast; S12 Fig, human). For each cluster, we selected a GO term with median size as the cluster representative. Among 45 clusters in yeast functional networks, only four, related to transcription (“ATP-dependent RNA helicase activity”, “RNA polymerase I activity”, “DNA-directed RNA polymerase II, core complex”, and “core TFIIH complex”) were biased towards both *k*- and *w*-functions (-log[*P*] ≥ 2, hypergeometric test). In contrast, 15 and 14 clusters were biased towards *k*- and *w*-functions, respectively. Clusters biased towards *k*-functions often represented cytokinesis (e.g., “septin complex”, “establishment or maintenance of actin cytoskeleton polarity”, and “mitotic cytokinesis”), while clusters biased towards *w*-functions corresponded to RNA degradation (e.g., “exosome [RNase complex]”, “ribonuclease MRP complex”, and “nuclear polyadenylation-dependent mRNA catabolic process”). Taken together, these results demonstrate that link clustering characterizes a unique subset of EGs with distinct biological functions. All *k*- and *w*-functions and their clusters in yeast and human were shown in S5 and S6 Table, respectively.

### EGs are clustered for the greater connectivity of *k*-dependent EGs

As we discussed in the Introduction, the clustering of EGs may largely be attributable to *k*-dependent EGs that are central, whereas *w*-dependent EGs particularly characterize non-central clusters with greater functional dependency. Indeed, we observed that there were more *k*-functions than *w*-functions (Fig 4A), indicating that *k*-dependent EGs are more likely clustered into a same function than *w*-dependent ones. To control the number difference between *k*- and *w*-dependent EGs, we randomly removed the same number of EGs and monitored the decrease of enriched GO terms. In the yeast consolidated network, for instance, we observed that removing *k*-dependent EGs lead to a greater decrease in the number of enriched GO terms compared to removing *w*-dependent ones for all three GO categories (Fig 4B). We observed similar results for the *k*- and *w*-dependent EGs from other PPI networks, with an exception of the yeast binary network (S13 Fig).

**Fig 4.**
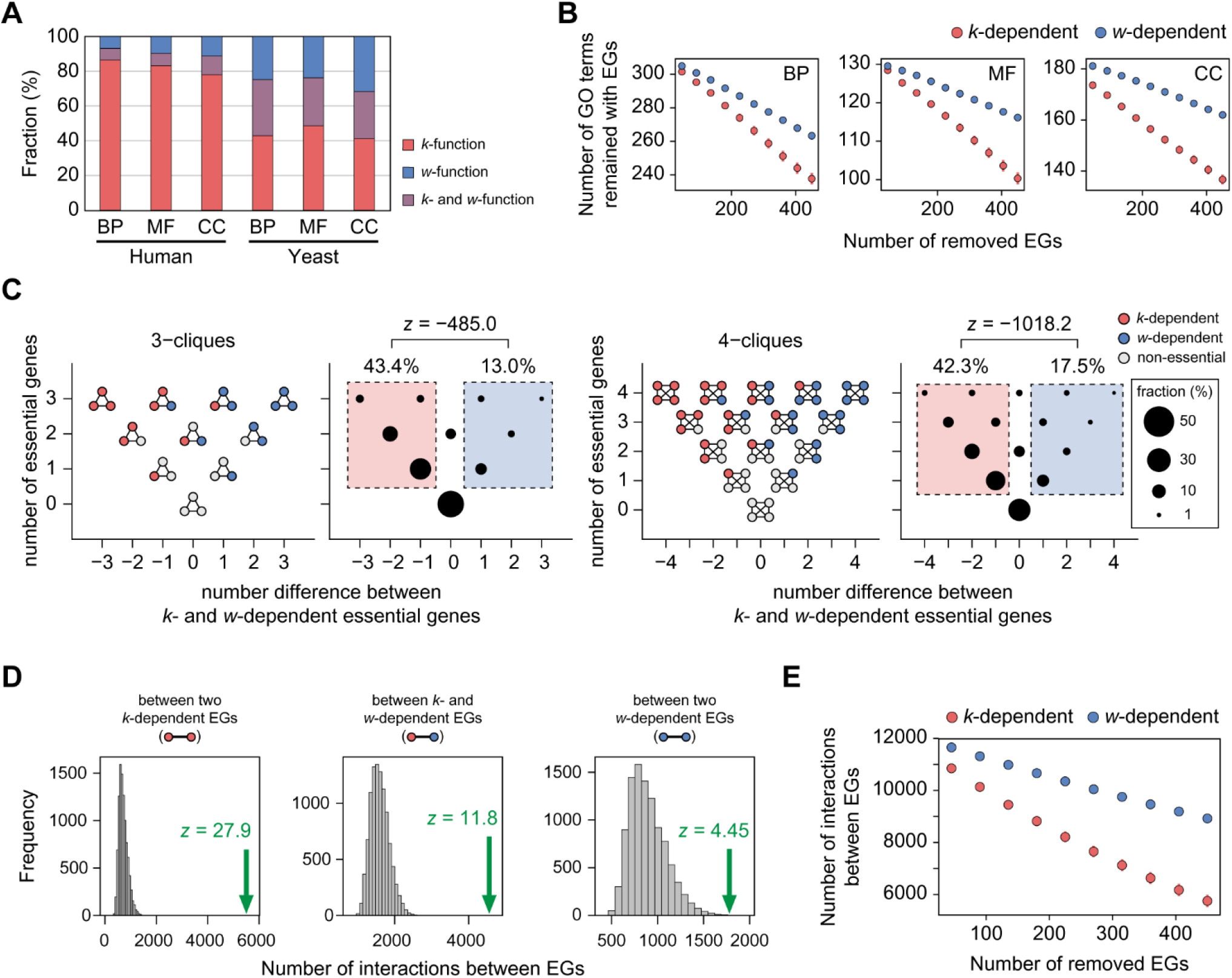
Contribution of *k*- and *w*-dependent EGs for the clustering of EGs. (A) Fraction of *k*- and *w*-functions in yeast and human for three GO categories. (B) Implication of *k*- and *w*-dependent EGs on the enrichment of EGs for GO terms. (C) n-cliques and their biases towards *k*- and *w*-dependent EGs. The normal distribution for the *z* value was approximated by binomial trials with success probability estimated by the fraction of *k*-dependent EGs. (D) Number of interaction between EGs (green arrow) and its comparison to random sets (grey bars), in which EGs and non-EGs were shuffled. (E) Implication of *k*- and *w*-dependent EGs on the number of interactions between EGs. The same number of *k*- and *w*-dependent EGs were removed randomly 100 times (error bar, 99% confidence interval).

To further examine clustered network structure around EGs, we explored *n*-cliques, which are fully-connected subgraphs with *n* nodes. In the yeast consolidated network, for instance, cliques frequently included more *k*-dependent EGs (Fig 4C; 3-cliques, fraction = 43.4%, 4-cliques, 42.3%) than *w*-dependent ones (3-cliques, 13.0%, *z* = −485.0; 4-cliques, 17.5%, *z* = −1018.2). We observed similar results in other PPI networks, with an exception of the human binary network (S14 and S15 Fig). This result indicates that the greater connectivity of *k*-dependent EGs likely give rise to the dense connectivity among EGs, hence their clustering.

It was also previously observed that EGs are frequently connected to each other in PPI networks [14, 15]. We found that interactions between EGs were more frequently found between *k*-dependent EGs, rather than *w*-dependent ones. In the yeast consolidated networks, for instance, the number of interactions between two *k*-dependent EGs (Fig 4D; *n* = 5,502) was greater than that between two *w*-dependent EGs (*n* = 1,780). By shuffling EGs and non-EGs, we observed that EGs were connected to each other more frequently than random expectation, with the greatest extent between two *k*-dependent EGs (z = 27.9), followed by a *k*-dependent EG and a *w*-dependent EG (*z* = 11.8), then by two *w*-dependent EGs (*z* = 4.5). To control the number difference between *k*- and *w*-dependent EGs, we randomly removed the same number of EGs and monitored the number decrease of interactions between EGs. We observed that removing *k*-dependent EGs exhibited to a greater decrease in the number of interaction between EGs compared to removing *w*-dependent ones. We observed similar results in other PPI networks (S16 and S17 Fig). These observations strongly indicate that the clustering of EGs is largely confounded by the greater connectivity of central EGs; and the link clustering with greater functional dependency characterizes a distinct subset of non-central EGs.

### *w*-dependent EGs significantly impact communities at low-level hierarchy

Next we examined how the clustering of *w*-dependent EGs is distinct from that of *k*-dependent ones. We hypothesized that *w*-dependent EGs would have greater impact on communities at low hierarchy compared with *k*-dependent ones, since such communities impose strong functional dependency among their members. To test this hypothesis, we investigated the impact of removing a node by monitoring changes in link density, Δ*D*, of communities. Because the level of hierarchy that would best illustrate such dependency among community members is unknown, we explored various hierarchical levels with varying *f*_H_, the fraction of proceeded merges between different communities; a larger *f*_H_ indicates a higher hierarchical level (see the Methods section).

Our studies revealed that impact on link density was greater with *w*-dependent EGs than *k*-dependent ones, and this difference was enhanced at lower hierarchical levels (Fig 5). For example, in the human consolidated network with *f*_H_ = 0.1, we observed a greater decrease in link density for *w*-dependent EGs (Δ*D*_w_ = −0.210; Fig 5A) than for *k*-dependent genes (Δ*D_k_* = −0.145) upon removal of a single node, suggesting that *w*-dependent EGs have a greater impact on community structure. In contrast, the difference in Δ*D* between *k*- and *w*-dependent EGs became extremely small at the highest hierarchical level (Δ*D_w_* - Δ*D_k_* = −0.00052, *f*_H_ = 1.0), suggesting that the effect of single node removal is unlikely to be distinguishable at the global network. Additionally, to confirm that Δ*D* is relevant to gene essentiality, we compared changes in link density upon the removal of a single node for non-EGs (Δ*D_n_*). At lower hierarchical levels (*f*_H_ ≤ 0.4), Δ*D_w_* - Δ*D_n_* < 0, indicating that *w*-dependent EGs have a greater impact on local community structure than non-EGs. Similar results were observed in other PPI networks: Δ*D_w_* - Δ*D_k_* < 0 (Fig 5B) and Δ*D_w_* - Δ*D_n_* < 0 (S18 Fig) at lower hierarchical levels with low *f*_H_ values.

**Fig 5.**
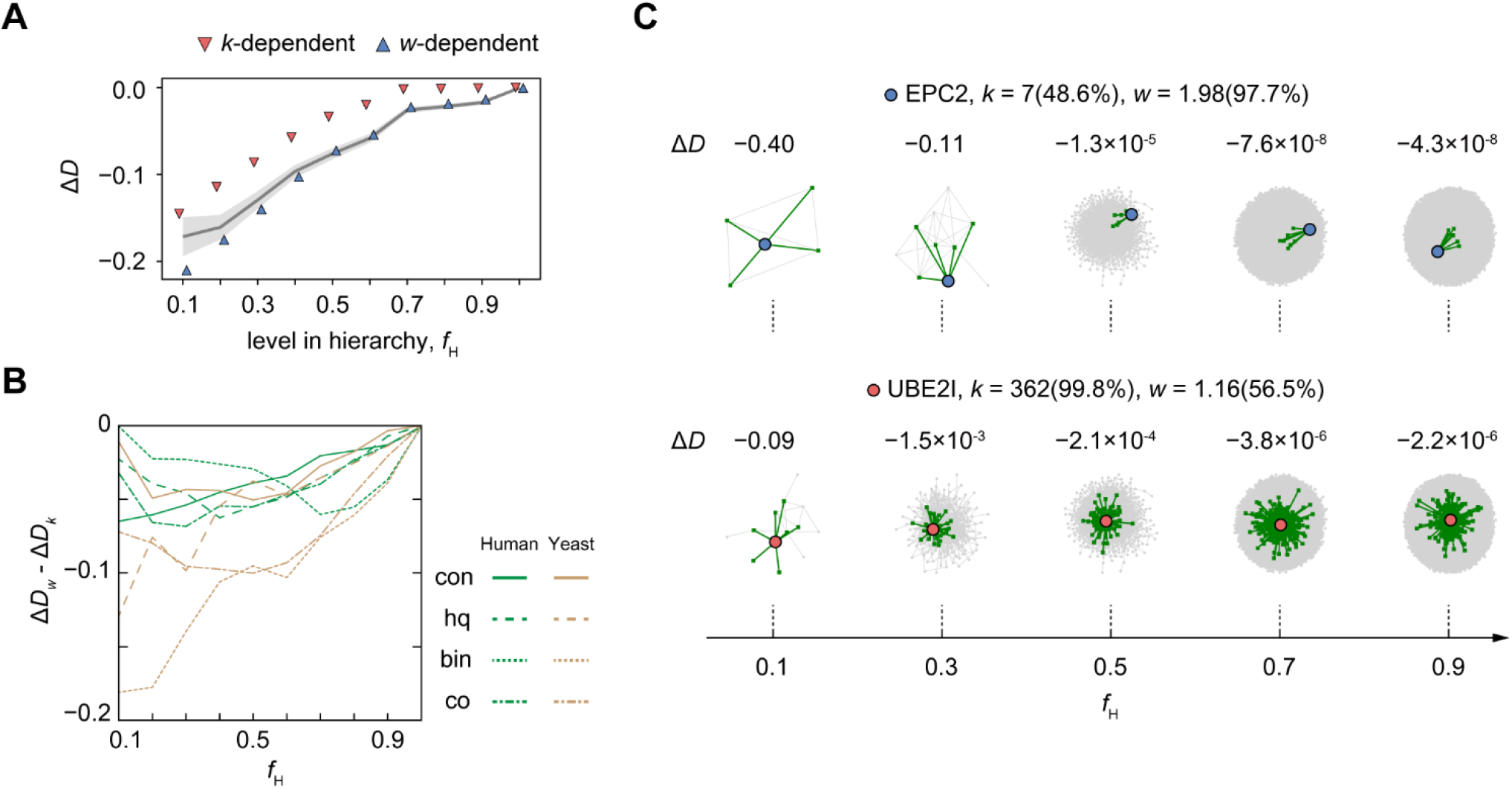
Impact of removing a single node on community structure with or without hierarchy. (A) Change in link density of the community structure upon removal of a single node (Δ*D*) for *k*- or *w*-dependent EGs at different hierarchical levels (*f*_H_). The grey line indicates the average Δ*D* for non-EGs (area, standard error). (B) Difference in Δ*D* between *k*- and *w*-dependent EGs in eight different PPI networks. Networks were designated as follows: con, consolidated; hq, high-quality; bin, binary; co, co-complex. (C) Examples illustrating a *w*-dependent essential gene *EPC2* and a *k*-dependent essential gene *UBE2I* for their impact on community structure. The protein of interest (blue circles, *EPC2;* red circles, *UBE2I*) and its interactions (green line) with its first neighbors (green circles) are shown in the community. Numbers in parentheses indicate the rank percentile for *k* and *w*.

An example of the impact of single node deletions on community structure with varying hierarchy is shown in Fig 5C. At *f*_H_ = 0.1, deletion of *EPC2*, a *w*-dependent essential gene with few and clustered links (*k* = 7 [rank percentile = 48.6%], *w* = 1.98 [97.7%]), had a large impact on the community structure (Δ*D* = −0.40), causing the removal of four of nine total links. In contrast, deletion of *UBE2I*, a *k*-dependent essential gene with many unclustered links (*k* = 362 [99.8%], *w* = 1.16 [56.5%]), had a smaller impact on the community structure (Δ*D* = −0.09), even though more links (*n* = 7) were removed. This observation illustrates how *w* can predict the impact of removing a single node on community structure at lower hierarchical levels. At higher hierarchical levels *f*_H_ ≥ 0.5), both *EPC2* and *UBE2I* are members of the same large communities; therefore, the impact of their deletion is much smaller (Δ*D*_EPC2_ = −1.3×10^−5^, Δ*D*_UBE2I_ = −2.1×10^−4^, at *f*_H_ = 0.5) than at lower levels of hierarchy. These results strongly indicate that a non-central EG’s functional dependency on its neighbors is crucial for its essentiality.

### Evolutionary histories of *k*- and *w*-dependent EGs are different

Given that *k*- and *w*-dependent EGs were distinct in terms of biological function (Fig 3) and network structure (Fig 5), it follows that they would also have different evolutionary trajectories. As EGs are indispensable by definition, they are more highly conserved than non-EGs [29, 30], but the difference seems not much strong [31]. Given that central genes tend to be well-conserved [32], *w*-dependent EGs may weaken the difference of conservation between EGs and non-EGs. To test this speculation, we investigated the extent of conservation of *k*- and *w*-dependent EGs using phyletic age and evolutionary rate. Briefly, the phyletic age of a gene is the most evolutionarily distant species group with homologues, and the evolutionary rate is defined as *d*_N_/*d*_S_, the ratio of nonsynonymous substitutions to synonymous substitutions.

We found that *k*-dependent EGs are older and evolve more slowly than *w*-dependent EGs. In the human consolidated network, orthologues of *k*-dependent EGs were significantly more biased towards distant species groups, such as “eukaryota” and “cellular organisms,” than were *w*-dependent orthologues (Fig 6A, *P* = 1.7×10^−41^; Mann-Whitney *U* test). Additionally, the average evolutionary rate of *k*-dependent EGs (Fig 6B, average *d*_N_/*d*_S_ = 0.222) was smaller than that of *w*-dependent EGs (average *d*_N_/*d*_S_ = 0.282), and the gene distributions were significantly different (*P* = 6.5×10^−24^). These results were replicated in most yeast and human PPI networks, except the binary networks (S19 and S20 Fig). Therefore, *k*-dependent EGs are conserved as expected, whereas *w*-dependent EGs are younger and evolve faster.

**Fig 6.**
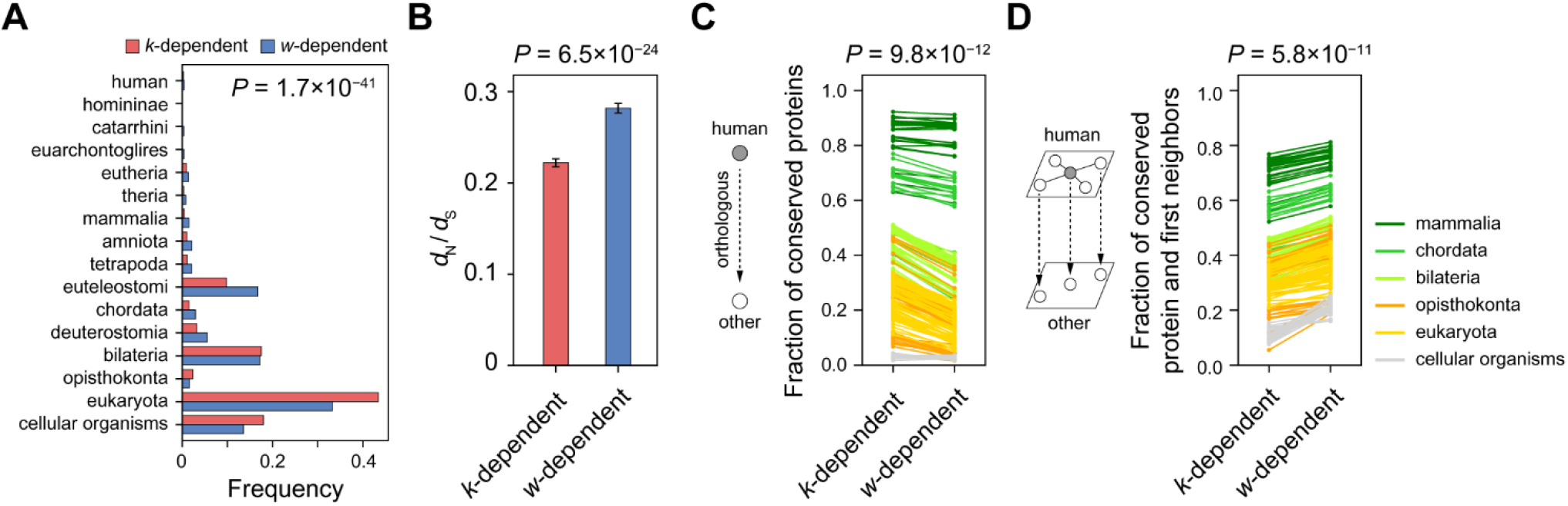
Conservation of *k*- and *w*-dependent EGs. (A) Phyletic age and (B) evolutionary rate of *k*- and *w*-dependent EGs. (C) Fraction of conserved proteins in 272 different species for *k*- and *w*-dependent EGs in the human consolidated network. The two fractions of *k*- and *w*-dependent sets are connected for each species. (D) Fraction of conserved members in groups consisting of a protein and its first neighbors for *k*- and *w*-dependent EGs.

Our hypothesis predicts that a *w*-dependent essential gene conserves together with neighbors, as its indispensability relies on the functional dependency with neighbors; in contrast, a *k*-dependent essential gene is expected to be conserved by itself. We indeed observed that, in the human consolidated network, *k*-dependent EGs exhibited a greater fraction of orthologues in other species than *w*-dependent EGs (Fig 6C, *P* = 9.8×10^−12^). In contrast, a *w*-dependent EG and its neighbors, considered as a group, showed a greater fraction of orthologues in other species compared with a group composed of a *k*-dependent EG and its neighbors (Fig 6D, *P* = 5.8×10^−11^). We observed similar results in other PPI networks, except the binary networks (S21 and S22 Fig). Taken together, these results show that *k*- and *w*-dependent EGs are distinct subsets of EGs, even with regard to the molecular evolution.

### Human *w*-dependent EGs are likely condition-specific

Growing evidences showed that gene essentiality is often not universal but specific to different conditions: a gene may change its essentiality across cell lines [7, 33–35] and species [36, 37]. Since central genes tend to be expressed broadly across cell lines and evolutionarily conserved, we may expect that *w*-dependent EGs, which are non-central, are specific to such conditions. To test this speculation, we investigated human EGs for their change of essentiality across different cell lines and between human and mouse.

We found that *w*-dependent EGs are more cell line-specific. We used a publicly available dataset of changing gene essentiality, which was tested across 10 different cell lines relying on large-scale targeted mutagenesis studies [7, 33–35]. In the human consolidated network, for instance, *w*-dependent EGs were biased toward smaller number of cell lines in which those genes were identified as essential, whereas *k*-dependent ones were skewed toward greater number of cell lines (Fig 7A; *P* = 1.2×10^−14^, chi-square test). Interestingly, gene essentiality is often specific to a single cell line, and we observed that *w*-dependent EGs were more often identified as essential in a single cell line (fraction = 35.3%) than *k*-dependent ones (26.6%). Moreover, across all the cell lines tested, *k*-dependent EGs were more often essential (18.0%) than *w*-dependent ones (8.5%). Other human PPI networks exhibited similar results (S23 Fig), with an exception of the binary network.

**Fig 7.**
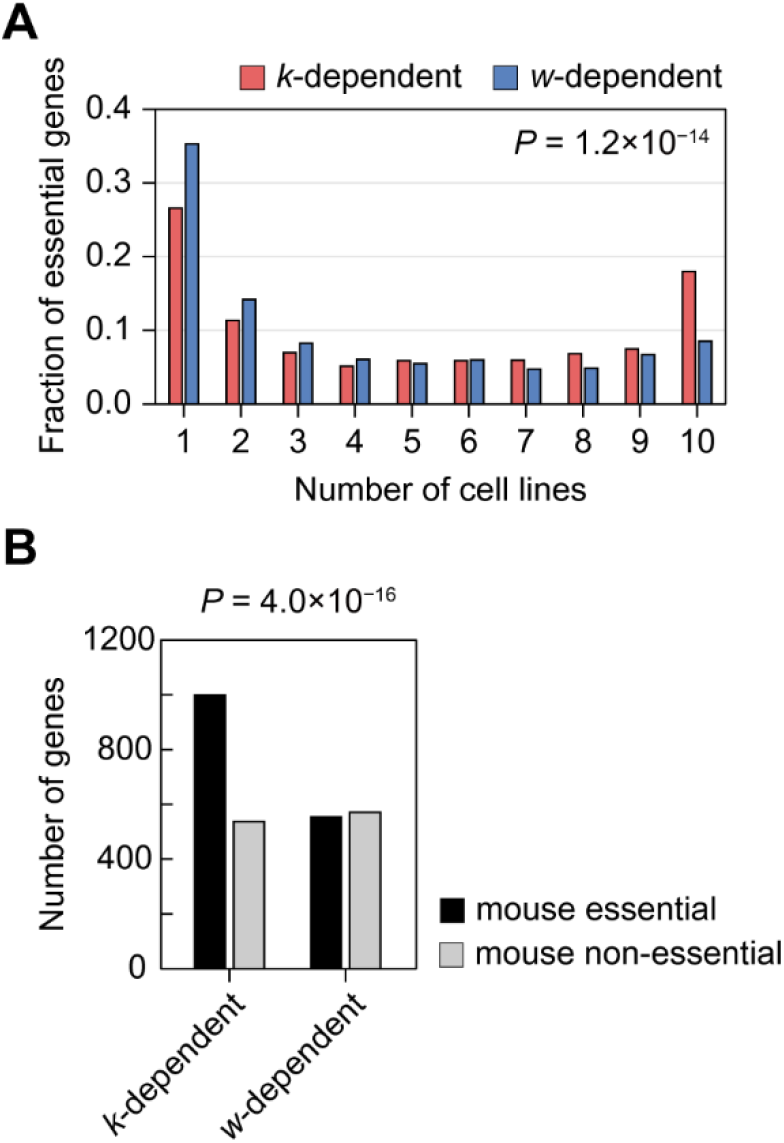
Change of essentiality across different cell lines and species in human *k*- and *w*-dependent EGs. (A) The distribution of *k*- and *w*-dependent EGs along with the number of cell lines in which given genes were identified as essential. (B) The essentiality of mouse orthologues of *k*- and *w*-dependent EGs.

We also found that *w*-dependent genes more frequently changed essentiality between human and mouse; that is, their essentiality is specific to human. In the human consolidated network, for instance, orthologues of *w*-dependent EGs were more frequently identified as non-essential in mouse (Fig 7B; fraction = 50.8%) than those of *k*-dependent ones (35.0%, *P* = 4.0×10^−16^, Fisher’s exact test), indicating essentiality of *w*-dependent genes were less evolutionarily conserved. We observed similar results in other human PPI networks (S24 Fig), with an exception of the binary network. Taken together, these results indicate that human *w*-dependent EGs are more condition-specific than *k*-dependent ones.

### Combining centrality with link clustering improves the prediction of gene essentiality

Considering that two network properties, centrality and link clustering, characterize distinct subsets of EGs, one may expect that utilizing both properties improves the prediction of gene essentiality than relying on a single property. Four centrality and two link clustering measures, and their combinations were assessed for the performance of gene essentiality predictions. Specifically, rank percentile of each measure and their additions were used as prediction scores, without any model-fitting or training parameters involved. As class imbalance is expected to be present (i.e., EGs are likely outnumbered by non-EGs), the recall-weighted average precision was taken as the performance measure [38].

We found that combinations of centrality and link clustering measures were more effective in the prediction of gene essentiality than those of two centrality or two link clustering measures. In human consolidated network, for instance, link clustering coefficient (LCC) was only the fifth in performance out of the six tested measures (Fig 8A, ‘LCC’). However, it achieved the first, the second, and the third in performance out of 15 measures, when LCC was combined with different centrality measures (Fig 8B; ‘C + LCC’, ‘E + LCC’, or ‘D + LCC’). By contrast, such improvement was not observed when two link clustering measures were combined (Fig 8B, ‘CC + LCC’). Indeed, we found that double-measure predictions utilizing both centrality and link clustering outperformed those relying only on two measures of a single property (Fig 8C, *P* = 6.0×10^−3^; Mann-Whitney *U* test). This observation was robust in different PPI networks; in 7 out of 8 PPI networks, the combination of centrality and link clustering measures held the best performance (S25 Fig) and exhibited significantly improved performances (S26 Fig).

**Fig 8.**
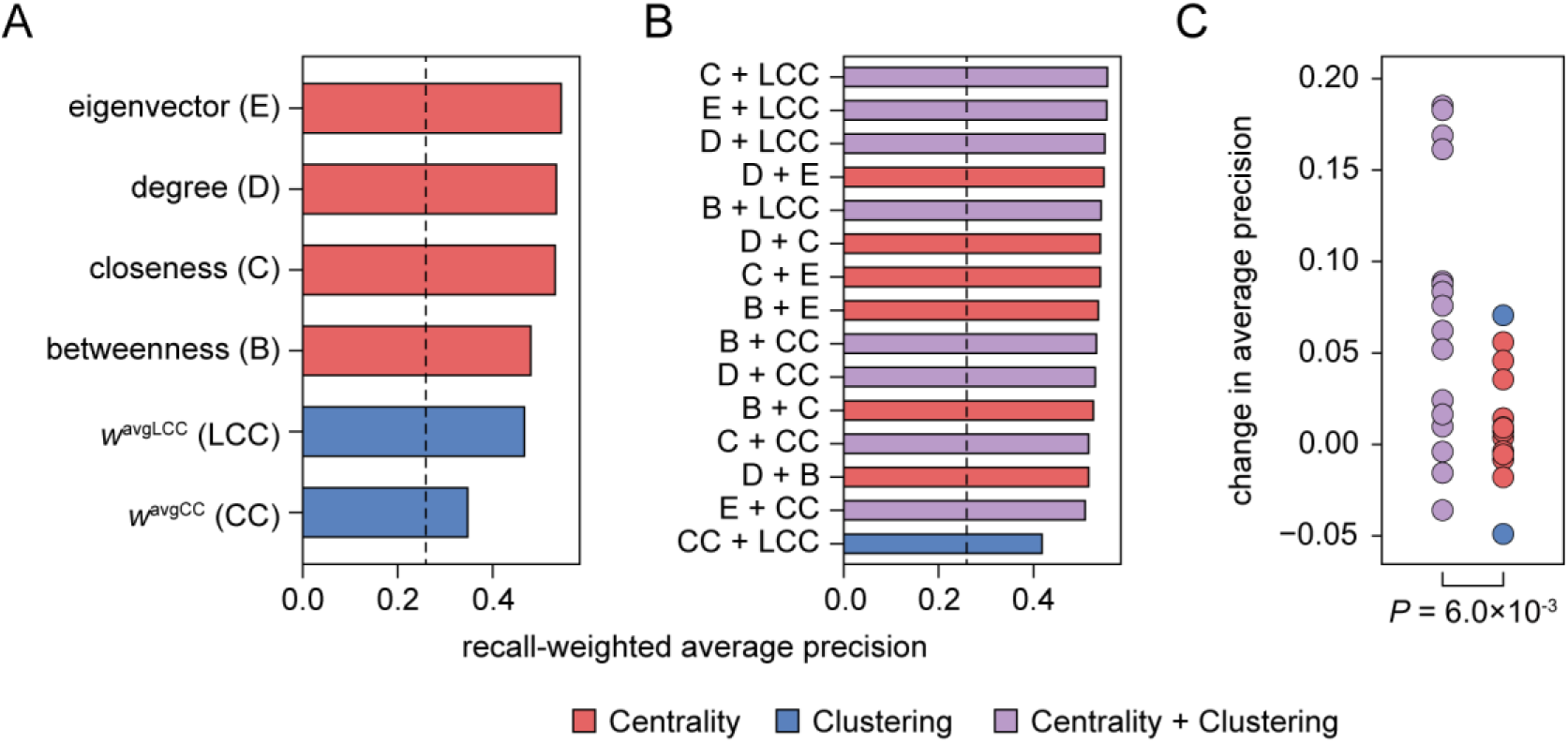
Combined effect of centrality and clustering measures to the prediction of essential genes. (**A**) Single-measure predictions. Rank percentile of a given measure was used as the prediction score. Precision-recall curve was explored and characterized by recall-weighted average precision. Dotted line indicates the expected precision by random sampling, which is the fraction of essential genes. (**B**) Double-measure predictions. Addition of the rank percentiles of two different measures was used as prediction score. (**C**) Changes in performance between double- and single-measure predictions.

## Discussion

In this study, we demonstrate that link clustering is capable of characterizing non-central EGs. Gene essentiality seems to be governed not by one, but by two distinct network properties: centrality and link clustering. Under what constraint cellular systems have evolved to confer gene essentiality with these two separate properties remains a matter of scientific exploration. As Dawkins stated, genes are “selfish” [39]; a gene may establish a large number of clustered links, rendering itself indispensable for the cooperative functions it partakes in and thus ensuring its persistence. However, this selfishness comes with spontaneous fitness costs to the population. Such cellular systems likely become more fragile by losing “error tolerance”, as random failure of non-central nodes may easily cascade globally throughout the greater dependency with central nodes. Therefore, selective pressure may have constrained cellular systems to have genes with either high centrality or high clustering, but not both. The results we present here support this hypothesis (Fig 9). Of note, this division of labor is further supported by the distinctive evolutionary modes of old and young EGs. Old EGs are known to be broadly conserved and confer commonly indispensable functions across species [29, 30]. In contrast, young EGs are likely involved in neofunctionalization for lineage-specific functions [36, 37]. Therefore, young EGs are less likely to be central genes, as the gain of new interactions (i.e., neofunctionalization) rarely occurs in a short period of time [40]. Indeed, we observed that *w*-dependent EGs are younger, fast-evolving, and less likely conserved across species than *k*-dependent EGs (Fig 6).

**Fig 9.**
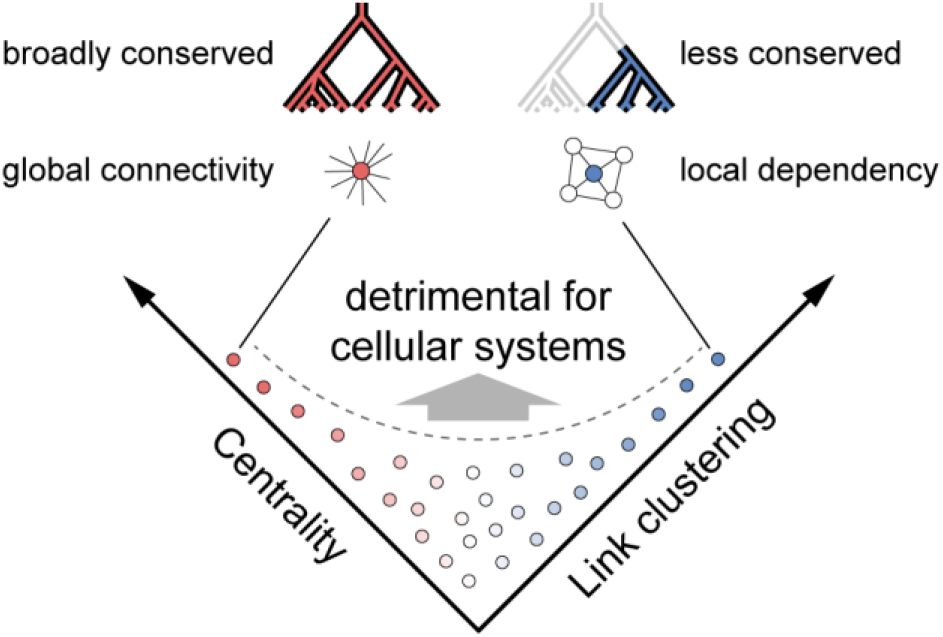
Division of molecular labor among essential genes and their characterization based on centrality and link clustering. Gene essentiality is indicated by an increase in either centrality or link clustering, shown as red or blue circles, respectively. Essential genes are defined by one of these two properties to avoid excessive deleterious impacts resulting from their combination.

Importantly, the discussion above asserts that links with strong dependency (“strong links”) are constrained from connecting to central nodes of greater importance in global connectivity; it also predicts that central nodes are connected by weak links. Indeed, several reports have shown such non-random distribution of link strength. In social networks, based on the duration of phone calls, strong links formed local communities, whereas weak links were more crucial for global integration [13]. In brain networks, strong co-activation of brain parts formed topological modules; therefore, efficient information flow at a global scale was achieved by introducing much weaker co-activation [41]. In genetic interaction networks, genes with highly correlated genetic interaction profiles formed densely connected modules, corresponding to specific pathways or protein complexes, whereas genes with low profile similarity corresponded to larger bioprocesses or cellular compartments at higher levels of hierarchy [42]. In PPI networks, interactions with balanced stoichiometry [27] or stable interface structure [43] often formed topological modules; in contrast, substoichiometric interactions had a greater impact on global network integration [27]. Therefore, as our hypothesis predicted, strong links with greater dependency are devoid of engagement with global connectivity in many real networks. This further supports our conclusion that functional dependency between nodes determines a link’s deleterious impact on its perturbation, and thus, gene essentiality.

Our investigation on network structure strongly suggests that functional dependency between nodes determines gene essentiality, rather than clustered structure itself (Fig 1D-G; Fig 4 and 5). Notably, different PPI networks describe distinct range of functional dependency between proteins, and those presenting weak dependency may delimit our analyses. For instance, Yu et al. showed that binary networks consist largely of transient signaling interactions and inter-complex connections [28], which likely convey lesser functional dependency than the interactions of other networks. Indeed, for the distinction of central and non-central EGs, binary networks often failed to behave as other networks do with respect to network structure (S4, S13, and S15 Fig) and biological characteristics (S20-24 Fig). In contrast, interaction proteomics experiments evaluate the consistency of physical binding between proteins, hence their dependency [24–27]. We found that the link clustering measures (*w*^avgCC^ and *w*^avgLCC^) were correlated with experimental link weights (*w*^avgEXP^) from those studies, and that experimental link weights were indeed capable of characterizing gene essentiality (S2 Fig). Moreover, we confirmed these observations in two recent unbiased human interactomes, BioPlex 2.0 [44] and HI-II-14 [45], which were detected by affinity-purification followed by mass-spectrometry (AP-MS) and yeast-2-hybrid (Y2H), respectively. In BioPlex 2.0, we observed that the link clustering measures (*w*^avgCC^ and *w*^avgLCC^) were correlated with gene essentiality (*f*_E_), but not with the degree (k) (S27 Fig). In contrast, in HI-II-14, link clustering measures were not correlated with *f*_E_, indicating that link clustering fails to characterize gene essentiality in this binary network. Taken together, link clustering is capable of characterizing non-central EGs in PPI networks encompassing strong dependency; and this further supports our hypothesis that functional dependency, rather than clustered structure itself, explains gene essentiality.

At a glance, the link clustering in this manuscript seems relevant to the fact that EGs are clustered, which was shown in many previous studies [16–21]; however, we posit that this is not the case. In our approach, the link clustering has been carefully tested for its orthogonality to centrality. However, those previous studies largely employed *a priori* knowledge of gene essentiality to define essential interactions and essential modules, which inherently include many central EGs. Indeed, without imposing functional dependency, we observed that *k*-dependent EGs are frequently clustered with and connected to each other than *w*-dependent ones (Fig 4). In addition, Song and Singh showed that EGs in hub modules satisfy modular behavior of EGs, as well as their high centrality [22]. Given that EGs are often clustered *and* central, those previous studies argued that central genes merely have greater chance to be involved essential interactions and modules, and thus centrality is not the cause of gene essentiality. Therefore, this argument focuses on central EGs and replaces the cause of their essentiality from centrality to biological interactions or modules, rather than separating non-central EGs. Note that we have neither an intention nor an evidence to argue whether or not network clustering is a real cause for a central gene to be essential. Instead, the link clustering in our approach clearly separates non-central EGs from others (i.e., central EGs and non-EGs), and in such case centrality remains as a partial contributor for gene essentiality.

Toward our goal for understanding non-central EGs, it seems crucial to properly aggregate the clustering of multiple links onto a single node, and here we have taken *the average* of link clustering. Interestingly, we noted that two previous studies have shown that *the sum* of link clustering also has ability to predict gene essentiality [46, 47]. By intuition, the sum would result in another centrality measure for its inherent correlation with the number of interactions, or the degree (*k*). We thus examined the difference between the average and the sum of link clustering measures (*w*^CC^, *w*^LCC^, and *w*^ECC^), with regard to their correlation with gene essentiality *f*_E_) and centrality measures (*k, BC, CC*, and *EC*). We found that the sum could characterize central EGs, unlike that the average characterized non-central ones. Despite that the sum correlated with *f*_E_, it often exhibited strong correlation with centrality measures (S2 Table), which was significantly greater than the average (S28 Fig). This indicates that the link clustering measures we present here provide a unique perspective on non-central EGs through a careful assessment of their orthogonality to centrality.

We observed that *w*-dependent EGs change their essentiality between human and mouse more frequently than *k*-dependent ones (Fig 7B). Since *w*-dependent EGs were determined by their greater link clustering, one might ask how the essentiality change is related with interacting neighbors. Interestingly, several hypotheses had been proposed for the essentiality change of functionally relevant genes across species. The “flipping” hypothesis explains that genes within a same protein complex flip together their essentiality between two species, depending on whether the function is essential in given species [20], whereas the “compensation” hypothesis is that one gene takes over the essentiality of other relevant gene through the replacement of the essential function [48]. Meanwhile, the “involvement” hypothesis is that genes often gain essentiality through being connected to conserved essential functions, leading to the lineage-specific divergence of gene essentiality [36]. We examined these evolutionary scenarios in our dataset by investigating the essentiality changes of *w*-dependent EGs with their neighbors. Specifically, given a *w*-dependent EG that is not essential in mouse, it follows i) the flipping hypothesis when its neighbor is also essential only in human, ii) the compensation hypothesis when its neighbor is essential only in mouse, or iii) the involvement hypothesis when its neighbor is essential in both human and mouse (S29 Fig). We found that *w*-dependent EGs prefer the involvement hypothesis over other hypotheses. Specifically, taking random sets by shuffling mouse gene essentiality, we observed that only the cases conforming to the involvement hypothesis were more frequent than random expectation (S29A Fig). However, we also noted that a vast majority of neighbors of *w*-dependent EGs did not follow any of the three evolutionary hypotheses (S29B Fig), indicating that it remains an open question how interacting neighbors had contributed to the changes of gene essentiality during evolution. We believe that our findings here may provide useful hints to a systematic investigation of the gene essentiality changes, in particular featuring the neighbors with clustered links.

Identification of cell line-specific EGs is important for finding therapeutic targets of diseases [49]. For instance, by targeting such genes, tumor cells can be selectively eliminated without excessive damage to normal cells [34, 35]. While genome-wide genetics studies directly identify the specificity of gene essentiality across cell lines, which is in particular accelerated by clustered regularly interspaced short palindromic repeat (CRISPR) system [50], it would be also insightful to characterize such genes. In recent studies, genes that are universally essential were shown to be conventional: they have many interaction partners and are highly expressed [7, 34]. In contrast, properties of cell line-specific EGs have remain largely illusive. Notably, we showed that *w*-dependent EGs were more cell line-specific (Fig 7), suggesting that functional dependency may underlie the varying essentiality of non-central genes across different cell lines. Therefore, we anticipate that network structure may provide a useful framework for understanding gene essentiality varying on different conditions.

## Methods

### Relationships between gene essentiality and network properties

The “consolidated” PPI networks were downloaded from the web interface to the Interaction Reference Index repository (iRefWeb) [51] on June 7, 2017. The “binary” and “co-complex” networks were downloaded from the high-quality interactomes (HINT) database [52] on June 27, 2017. The “high-quality” networks were created by combining the “binary” and “co-complex” networks. Gene essentiality information was downloaded from the online gene essentiality (OGEE) database [53] on June 9, 2017. Any essentiality annotation with the “TextMining” data type was removed.

To explore a parameter’s ability to characterize gene essentiality, we calculated the Pearson correlation coefficient (*R*) between the fraction of EGs (*f*_E_) and the average of given parameter along with the rank-ordered groups. Proteins were sorted based on increasing order of the parameter of interest and added into a single bin until the bin contained at least 2% of the total population. This applied to all relationships of *f*_E_ with centrality measures (*k*[*DC*], *BC, CC*, and *EC*), clustering measures (*C, w*^CC^, *w*^LCC^, and *w*^ECC^), and experimental link weights (*w*^EXP^). The experimental link weights from Hein et al. [27] was scaled in log10.

For the linear curve fitting between 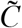 and *w*^avgCC^, we used the linalg.lstsq() function in the Python “numpy” package; optimal *β* was found for a linear curve *y* = *β_x_* + *α*, where *y* = *w*^avgCC^ - 1, 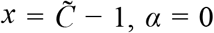. The *p*-value of the difference between *β*_E_ and *β*_NE_ was transformed from a normal statistic *z* = (*β*_NE_ - *β*_NE_) / (var[*β*_E_] + var[*β*_NE_])^1/2^, where var(*β*) = *σ*^2^/*ns*^2,^ *σ* is the residual variance, and *s* is the sampled variance of *x* [54].

For the principal component analysis of EGs, we used decomposition.PCA() object in the Python “scikit-learn” package. To scale the features, we also used transform() function of preprocessing.StandardScaler() object in the same package.

### Monitoring global and local connectivity upon pruning

Pruning analysis was performed in a manner similar to that previously reported [11]. Proteins were progressively removed from a given network at 5% of the total protein population in decreasing order of *k* and *w*, monitoring corresponding changes in *R*_GC_ and Δ*C* with varying *f* (the fraction of removed proteins). Random sets for *r*-pruning were constructed by shuffling the rank-ordered proteins 100 times. To calculate the Δ*C* of individual nodes, we constructed 100 random networks by degree sequence preserved randomization [11, 55] and subtracted the mean node clustering coefficients of random sets from the observed node clustering coefficient.

For simplicity, we here explain about *k* and *w*, but the same analysis was applied for all centrality and clustering measures used. Differences in the impact of *k*- and *w*-pruning over *r*-pruning were quantified by comparing the area under the curve for various parameters. As Δ*C* showed varying scales in different PPI networks, we normalized the monitoring parameters to range from 0 to 1, as follows: *x*_norm_ = (*x*_obs_ - min[X])/(max[X] - min[X]), where x represents a monitoring parameter (*R*_GC_ or Δ*C*), xobs represents the observed value, and X represents a set of observed values from all pruning analyses (*k*-, *w*- and *r*-pruning). We then calculated the trapezoidal area over *f* = [0, 0.95].

### Classification of *k*- and *w*-dependent EGs

Here a gene can only be classified one of two ways: essential or non-essential. We assigned a value of 1 to EGs, and 0 to non-EGs. The probability that a given gene is essential was then calculated using logistic regression analysis according to a leave-one-out scheme, with *k* and *w* as dependent variables, resulting in *P*_E_(*k*) and *P*_E_(*w*), respectively. We performed the logistic regression analysis using the Python “scikit-learn” package. In addition, k_c_ (w_c_) was determined to maximize MCC regarding all nodes with *k* ≥ k_c_ (*w* ≥ w_c_) as predictive positives.

### Functional association between *k*- and *w*-dependent EGs

To construct functional networks, we defined GO terms as *k*-functions for those enriched with *k*-dependent genes in at least three PPI networks (*P* < 0.05, hypergeometric test); *w*-functions were defined accordingly. GO terms were discarded when the number of annotated gene was less than three. Note that a GO term can be enriched with both *k*- and *w*-dependent EGs, as their enrichment was tested independently. A link was established between two GO terms if there was significant gene overlap (*P* < 10^−5^) between the two GO terms. We used the MCODE application [56] to identify clusters in the functional networks. For each cluster, we selected the median-sized GO term as the cluster’s representative function, where size is the number of genes in that function. We constructed in total six functional networks for three GO categories (BP, MF, and CC) and two eukaryotic species (yeast and human). Annotations were downloaded from the GO database [57, 58]; the submission date of the human data used in the study was September 26, 2017, and that of yeast data was September 13, 2017.

### Impact of removing a node on community structure

For each PPI network, we constructed a hierarchical organization based on the *Walktrap* algorithm, using the Python package “python-igraph”. The algorithm relies on random walk to measure the similarity between two nodes by comparing their probability of random visits on other nodes: if two nodes are in a same community, the random walk starting from each of the two nodes will visit all the other nodes in the same way [59]. Established the similarity between nodes, the clustering process is agglomerative; in earlier steps nodes with greater similarity are put together into a community. Therefore, we took the fraction of passed merge steps, *f*_H_, as an indication of hierarchy; the smaller the *f*_H_, the lower the hierarchical level. Increasing *f*_H_ by 0.1, communities at different levels in the hierarchical organization were then collected. Communities comprising less than three members were discarded. The change in link density in community *s* upon the deletion of node *i* was calculated as follows: Δ*D_s,i_* = (*l_s_* - *l_s,Δi_*)/(*n_s_*×(*n_s_* - 1)/2), where *l_s_* represents the number of links within *s* (i.e., two end nodes are both the members of *s*), *l_s,Δi_* represents the number of links within *s* after removing node *i*, and *n_s_* represents the number of members in *s*. Therefore, Δ*D* represents the proportion of removed links upon a node deletion, indicating the extent of functional dependency within a community relying on that node.

### Conservation of EGs and their partners

We gathered orthologues of human (or yeast) proteins among 272 species from the Inparanoid database (version 8.0) [60]. We then assessed the conservation of *k*- and *w*-dependent EGs in two different ways: based on individual proteins (*CSV*_P_) or based on proteins and their interaction partners (*CSV*_NB_). For a species *x, CSV*_P_ = Σ_*i*_ O(*i,x*)/Σ_*i*_, where O(*i,x*) = 0 if a protein *i* has no orthologue in species *x*, and O(*i,x*) = 1 otherwise. Additionally, *CSV*_NB_ = Σ_*i*_ [(1+Σ_*j*_ O[*j,x*])/(1+Σ_*j*_)]/Σ_*i*_, where *j* represents the first neighbors of *i*. To demonstrate that the results were unbiased toward any specific phyletic group, species in six phyletic groups were visualized based on the common tree obtained from the National Center for Biotechnology Information. Gene ages were downloaded from the ProteinHistorian database [61]; specifically, we used protein families predicted from the OrthoMCL and PANTHER databases, and ancestral history reconstructed by Dollo parsimony. We used the pre-calculated set of *d*_N_/*d*_S_ for yeast [62] and human [63], for which evolutionary rates were computed with several species and the average was taken.

### Prediction of gene essentiality

We tested four centrality measures (degree, betweenness, closeness, and eigenvector) and two link clustering measures (*w*^avgCC^ and *w*^avgLCC^) for their predictive capability of gene essentiality. For single-measure predictions, the link percentile of each measure was used as prediction score. For double-measure predictions, the addition of two rank percentiles was used as prediction score. To evaluate the performance, precision-recall curves were drawn and the average precision was calculated in recall-weighted manner as follows: average precision = Σ(*RC_i_* - *RC_i-1_*)*PR_i_*, where *PR_i_* = *TP_i_* / (*TP_i_* + *FP_i_*) and *RC_i_* = *TP_i_* / (*TP_i_* + *FN_i_*); *TP_i_, FP_i_*, and *FN_i_* is the number of true positives, false positives, and false negatives at *i*th operation.

### Context-specific gene essentiality

For the gene essentiality in different cell lines, we used the dataset compiled in Bertomeu et al. [7], which includes gene essentiality tested in three other targeted mutagenesis studies [33–35]. For the gene essentiality in mouse, we used OGEE database [53]; orthologous genes between human and mouse were identified by using Inparanoid database [60].

## Supporting Information

**S1 Fig. Least square fit curves between 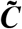 and *w*^avgCC^ for essential and non-EGs**. Slopes of least square fits for essential and non-EGs are shown as red and black dotted lines, respectively. The 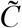 and *w*^avgCC^ of essential and non-EGs are shown as orange and grey circles, respectively.

**S2 Fig. Correlation of link weight from interaction proteomics experiments with gene essentiality and clustering measures**. Scatter plots illustrate *w*^avgEXP^ with (A) *f*_E_, (B) *w*^avgCC^, and (C) *w*^avgLCC^.

**S3 Fig. Correlation of clustering measures with *f*_E_ and other centrality measures, including betweenness, closeness, and eigenvector centrality**.

**S4 Fig. Principal component analysis on EGs in different PPI networks**. The variance explained by the components was shown in parenthesis. Centrality and clustering measures were shown in red and blue arrows, respectively.

**S5 Fig. Impacts of centrality and link clustering measures on global and local connectivity quantified by area difference between pruning curves**. Comparisons were made between four centrality measures: degree [*k*], betweenness, closeness, and eigenvector, and two clustering measures: (A) *w*^avgCC^ and (B) *w*^avgLCC^.

**S6 Fig. Pruning curves for *w* = *w*^avgCC^ and centrality measures in 8 PPI networks**.

**S7 Fig. Pruning curves for *w* = *w*^avgLCC^ and centrality measures in 8 PPI networks**.

**S8 Fig. Probability of a gene being essential based on *k* and *w* for all EGs and the criteria to classify *k*- and *w*-dependent EGs in eight different PPI networks**. Criteria are shown as follows: grey dotted lines, *P*_E_(*k*) = *P*_E_(*w*); red dotted lines, *k* = k_c_; and blue dotted lines, *w* = w_c_.

**S9 Fig. Finding the cutoffs, k_c_ and w_c_, to maximize MCC**. Red circles indicate points of maximum MCC. Numbers in parentheses indicate *k* or *w* with the maximum MCC.

**S10 Fig. Functional networks for different GO categories**. (A) The number of GO terms connected to the largest connected component in the functional network. (B – F) Visualization of other functional networks.

**S11 Fig. Number of links between *k*- and *w*-functions in real function networks (green arrow) and in random sets (grey bars)**. Random sets were constructed by shuffling *k*- and *w*-function tags 10.000 times.

**S12 Fig. Representative GO terms of clusters from human functional networks and the bias of clusters towards *k*- and *w*-functions**.

**S13 Fig. The implication of *k*- and *w*-dependent genes on the modular behavior of EGs**. The number of GO terms enriched with *k*- or *w*-dependent EGs were monitored after removing the same number of *k*- and *w*-dependent EGs, which were randomly selected 100 times.

**S14 Fig. Biases in 3-cliques towards *k*- and *w*-dependent EGs in different PPI networks**.

**S15 Fig. Biases in 4-cliques towards *k*- and *w*-dependent EGs in different PPI networks**.

**S16 Fig. Number of interaction between EGs (green arrow) and its comparison to random sets (grey bars) in different PPI networks**. Random sets were constructed by shuffling EGs and non-EGs 10.000 times.

**S17 Fig. The impact of *k*- and *w*-dependent genes on essential links**. Essential links were defined as interactions between EGs. The number of such interactions were monitored after removing the same number of *k*- and *w*-dependent EGs, which were randomly selected 100 times. Error bars indicate 99% confidence interval.

**S18 Fig. Change in link density of the community structure upon removal of a single node (ΔD) at different levels of hierarchy for the eight different PPI networks examined**. The average changes in link density for *k*- and *w*-dependent EGs are shown as red downward and blue upward triangles, respectively. Grey lines indicate the averages for non-EGs (area; standard error).

**S19 Fig. Phyletic age of *k*- and *w*-dependent EGs in different PPI networks**.

**S20 Fig. Evolutionary rate (dN/dS) of *k*- and *w*-dependent EGs in different PPI networks**.

**S21 Fig. Fraction of conserved proteins in different PPI networks**.

**S22 Fig. Fraction of conserved members in groups consisting of a protein and its first neighbors for *k*- and *w*-dependent EGs in different PPI networks**.

**S23 Fig. The distribution of *k*- and *w*-dependent EGs from different PPI networks along with the number of cell lines**.

**S24 Fig. The essentiality of mouse orthologues of *k*- and *w*-dependent EGs from different PPI networks**.

**S25 Fig. Performance comparisons of centrality and link clustering measures in 8 different PPI networks**.

**S26 Fig. Change in performance between double- and single-measure predictions in 8 different PPI networks**.

**S27 Fig. Correlation of link clustering measures (*w*^avgCC^ and *w*^avgLCC^) in BioPlex 2.0 and HI-II-14 with the fraction of essential genes (*f*_E_) and interaction degree (*k*)**.

**S28 Fig. Comparison between the average and the sum of link clustering measures**. Grey and black dotted lines show the mean observed correlation of the average and the sum, respectively, with centrality measures among different PPI networks (*P*, t-test).

**S29 Fig. Evolutionary hypotheses of essentiality change of *w*-dependent EGs and their neighbors**. (A) Number of neighbors following given hypotheses. Random sets were constructed by shuffling mouse gene essentiality, after determining the essentiality change of *w*-dependent EGs with real data (*n* = 10,000). (B) Fraction of neighbors following given hypothesis in all the neighbors of *w*-dependent EGs.

**S1 Table. Properties of PPI networks**.

**S2 Table. Correlation between gene essentiality and centrality and clustering measures in PPI networks**.

**S3 Table. Yeast *k*- and *w*-dependent essential genes**.

**S4 Table. Human *k*- and *w*-dependent essential genes**.

**S5 Table. Yeast *k*- and *w*-functions and theri clusters in function networks**.

**S6 Table. Human *k*- and *w*-functions and theri clusters in function networks**.

